# Numeric Analysis of Temperature Distribution in Man using a 3D Human Model

**DOI:** 10.1101/330357

**Authors:** Sipho Mfolozi, Arnaud Malan, Tunde Bello-Ochende, Lorna J. Martin

## Abstract

Premortem three-dimensional body temperature is the basis on which post-mortem cooling commences. Thermo-numeric analysis of post-mortem cooling for death-time calculation applies pre-mortem three-dimensional body temperature as *initial conditions*; therefore, an accurate determination of this distribution is important. To date, such prediction is not performed. This paper presents a thermo-numeric analysis method of predicting premortem three-dimensional body temperature in man, to be applied in thermo-numeric analysis of the post-mortem interval using the finite-difference time-domain method. The method applied a Pennes BioHeat Equation modified to linearize organ metabolic and blood flow rates with temperature in a transient thermo-numeric analysis scheme to predict naked three-dimensional temperatures of an MRI-built, 3D human model having 247 segmented organs and 58 categories of material properties under chosen *boundary conditions*. Organ metabolic heat and blood perfusion rates appropriate for a chosen pre-mortem physical activity, and known organ physical and thermal properties, were assigned to each organ. A steady-state temperature equilibration occurred after 8400 seconds. Predicted organ temperatures were topographically inhomogeneous. Skin temperatures varied between 20.5°C and 42.5°C, liver capsule temperatures were lower than parenchymal, and rectal luminal temperature were uniform.

## Introduction

Accurate death-time calculation facilitates medico-legal death investigation and ultimately improves the effective administration of justice. Thermometric death-time calculation methods are based on time-dependent temperature changes that occur in the body from a person’s time of death. Such methods match post-mortem temperature measurements with time-temperature curves modelled either empirically or numerically. The former are predetermined and valid for cooling conditions and thermometry techniques similar to those of the experiment. For example, deep rectal temperatures should be applied to the rectal temperature nomogram method [1]. The method cannot, therefore, be applied in non-predictable cooling conditions such as in the presence of strong thermal radiation. Numerically modelled methods are based on physical laws of heat transfer and are applied on geometries representing the human body using computational analyses under similar conditions. Numerical methods can theoretically be applied for any cooling condition, that is, if all the parameters required for calculation are available.

Both empiric and numeric current methods are formulated on the basis that the core temperature at death is ±37°C. Numerical methods require initial three-dimensional temperature at death, and the skin temperature values often applied are of a naked man at rest in a thermo-neutral zone, i.e. at an ambient temperature that does not elicit thermoregulatory changes in metabolic heat generation or evaporative heat loss. Core and skin temperatures in empiric methods are often assumed to be 37°C and 27°C, respectively. The Finite Element Model [2] linearly approximated the temperature gradient between a core of 37°C and skin of 27°C. This paper presents a method of explicit calculation of core, skin, and total-body temperatures present in a body at the time of death. Human surrogates on which post-mortem cooling research was historically undertaken include recently-deceased human bodies [3-5], cooling dummies [1, 6, 7], a wooden cylinder [8], the Finite-Element model, and most recently [9] a CT-based finite element model. The anatomical morphology of a wooden cylinder or cooling dummy is less than ideal compared to a human body: there were no individual organs, and just one computational domain is regarded as being representative of the average human tissue. The anatomical morphology of the Finite-Element Model was humanoid, and consisted of several, broad organ-categories that predominantly belong to anatomic regions. This study applied a 3D human model built from magnetic resonance images (MRI) of a living and healthy adult as a human surrogate. The model consisted of individually segmented and realistically shaped internal organs. The hypothesis is that predicted premortem temperature using such a model would be as close to reality as possible for purposes of death-time calculation, all else being equal.

In calculating the death-time interval, the Finite Element Model applied a transient thermal analysis throughout, from the time the model was alive until it had cooled to ambient temperature. This was done using a combination of heat density proportionally divided by organ volume, as well as by using a calibrated decrease-rate of internal power production to mark metabolic cessation at death. In contrast, this study predicts total-body temperature distribution in life as a separate process, so that calculation of post-mortem cooling would be a follow-on process. The analysis solution from the prediction can then applied as initial conditions of a follow-on transient thermal analysis that simulates post-mortem cooling only, in which organ metabolic heat and blood flow rates are set to zero. A separation of the two processes clarifies issues and makes for easier workflow. The topic of the follow-on transient thermal analysis for predicting post-mortem cooling may be the subject for a future paper.

Total-body temperature distribution in the pre-mortem period results from complex interactions of metabolic heat generation, which itself depends on multiple factors including physical activity, physiology and pathology states, as well as ambient temperatures, clothing/skin insulation, and prevailing thermoregulatory responses. Ideally, all these variables should be included in thermonumeric analyses of premortem total-body distribution. The method presented in this paper applied only endogenous metabolic heat and blood flow values of individual organs appropriate for a given pre-mortem physical activity from the literature. Numerical modelling of the human thermoregulatory system to predict core, skin, and total-body temperature distribution under various environmental conditions is applied in many fields of science such as heating, ventilation, and air-conditioning engineering (HVAC), thermal comfort, sports medicine, automotive, and aerospace industries. The first quantitative relationship that described heat transfer in living human tissue, which included the effects of blood perfusion on tissue temperature on a continuum basis, was presented by Pennes in 1948 [10]. His bio-heat equation (PBE) calculated the transient temperature distribution of the human forearm. The model included radial conduction of heat of the cylindrical forearm, metabolic heat generation, convection of heat by circulating blood, and heat loss from skin surface by thermal convection, thermal radiation, and thermal conduction. The Pennes bio-heat equation is expressed as:

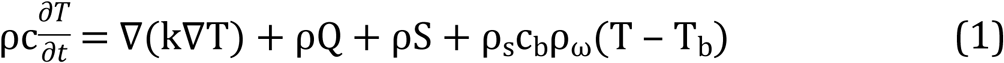

where ρ is organ density (kg/m^3^), c is organ specific heat capacity (J/(kg.K)), T is organ temperature (°C), t is time (s), k is its thermal conductivity (W/(m.K)), Q the organ’s specific metabolic heat generation rate (W), S is the organ’s specific absorption rate, and the subscript b denotes blood. Generated heat, furthermore, is spread by thermal diffusion. In the equation, metabolic body processes constitute a continuous heat source, which is specific to each tissue and assumed to be homogeneous within each individual tissue domain. Similarly, blood circulation acts as a homogeneous, tissue-specific heat-sink term. The PBE has been cited no less than 3720 times in research involving bio-heat transfer.

A number of bio-heat models were developed subsequent to the PBE [11 – 18], and many were incremental improvements addressing specific shortcomings. In all these models the morphology of the human body were schematic rather than anatomical, consisting of cylindrical segments representing hands, arms, forearms, head, neck, chest, abdomen, pelvis, thighs, legs, and feet. Each segment generally consisted of concentric layers of bone in the core followed by muscle, fat and finally skin on the surface. These models became more complex over time, incorporating active thermal control systems (metabolic, blood circulation & vasomotor) and passive thermal control systems (sensors, sensory data-processing). Functional relationships between organs become detailed and increasingly more accurate. Over the years, the anatomic morphology of their human surrogates evolved to realistic topography using 3D laser scanning. Some models could even sweat under the appropriate conditions. These models were used to predict skin temperature, thermal comfort and physiological responses to external thermal stimuli under various conditions. They were not used for and are unsuitable for predicting premortem total-body temperature distribution due to lack of realistic internal organs. This is the first paper to predict pre-mortem total-body temperature distribution.

## Materials

- Computer Hardware Analyses were carried out in a 2015-model Apple Mac Pro computer with a 12-core 3.5GHz processor and a 64GB RAM, which ran 64-bit Windows 8.1. Analysis data were stored in a 10TB external hard drive.
- Thermal Analysis Software A thermodynamic solver is computer software that solves heat transfer problems involving specified geometries based on physical laws that govern heat flow by thermal conduction, thermal convection, and thermal radiation. Such geometries may be a novel prototype or an untested design that is either too expensive to build or too complex for detailed laboratory analysis. Many thermal solvers are commercially available for licensed application. The thermodynamic solver used in this study was P-Thermal® and was an integral part of a larger biophysics simulation platform called Sim4Life® (IT’IS Foundation, Switzerland). The P-Thermal® solver was based on Poisson differential equation and enabled for the modelling of heat transfer in living tissue with a set of flexible boundary conditions. Some of P-Thermal® applications are in radio frequency hyperthermia, radio frequency tumor ablation, design and the optimization of ultrasonic devices, cryosurgery, hypothermia, pacemaker-implants, thermal effects of mobile phone use, temperature impact on neuronal dynamics, etc. The transient thermal solver assumes that a transient state exists, which requires all tissue domains to have non-zero thermal conductivity or nonzero heat transfer rates. Sim4Life® version 3.4.1.2244 was used in this study.
- 3D Human Model Our 3D human model was part of the Virtual Population 3.0––a set of fifteen high end, high resolution, whole-body computational human avatars built from MRIs of healthy male and female, adult and children volunteers [19]. The realistic shape, relative size, and anatomical relations of their organs were anticipated to afford realistic analysis of heat transfer and temperature distribution that would otherwise not be possible with concentric-style human models used in HVAC studies that historically applied the PBE. The 3D human model chosen for this study was Duke V3.0b03: a 1.77m tall and 72.4kg, 34-year-old male consisting of 247 computational tissue domains belonging to 58 tissue-property categories (Fig 1). All of Duke’s computational domains could be highlighted and manipulated individually in the thermal solver software.

**Fig. 1.**
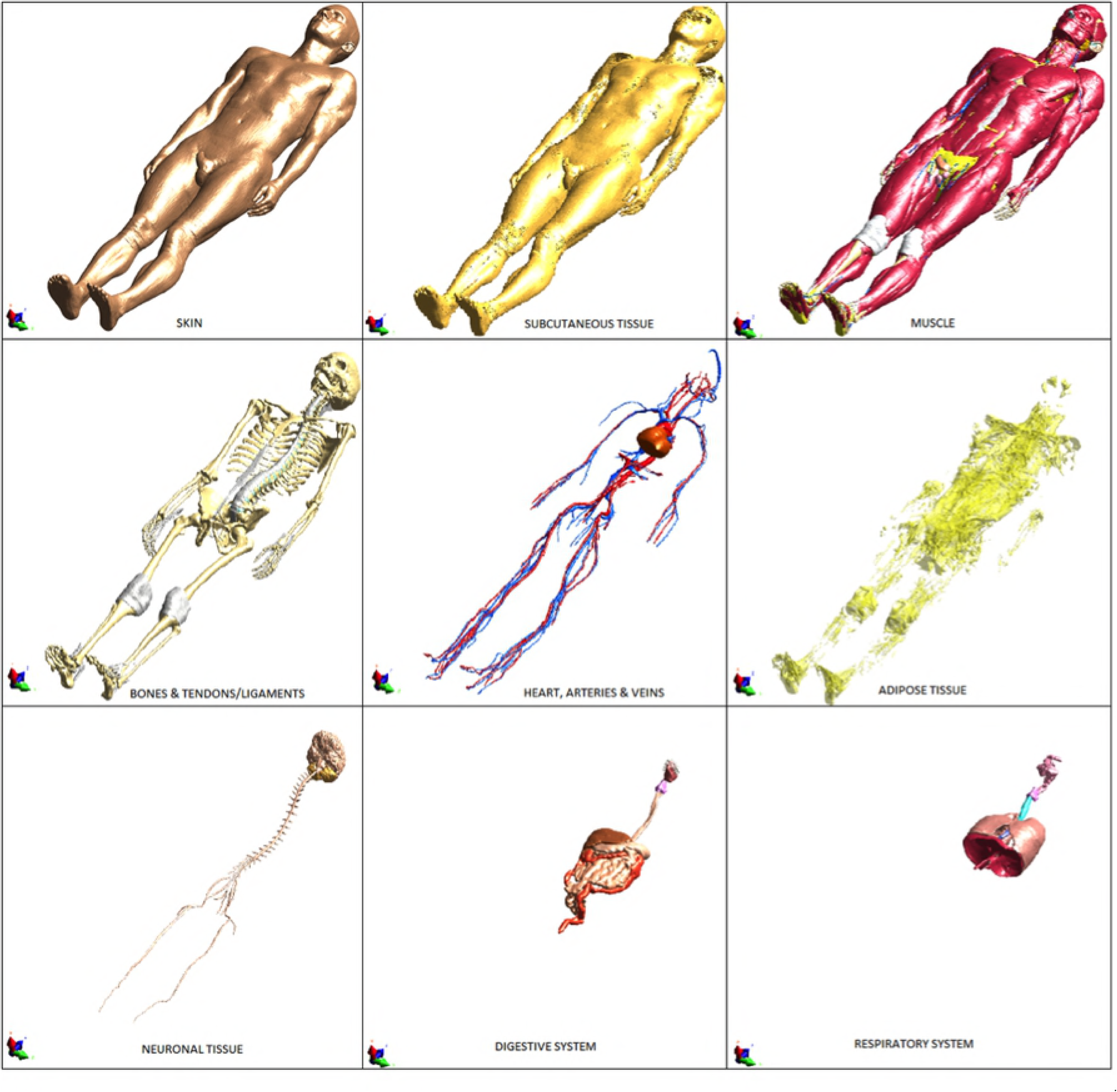
The 3D male model Duke v3.0b03 segmented into different systems. Top row from left to right shows skin, subcutaneous tissue and muscle. Middle row from left to right shows bones and tendons, the cardiovascular system and adipose tissue. Lower row from left to right shows the neuronal tissue, digestive system and the respiratory system.

The substantial number of tissue domains was due to several factors. Computational tissue domains were identified by anatomical name as identified by the MRI. Many tissue domains, such as the model’s bones, shared identical tissue properties. Some tissue domains were ‘non-human’ in the biological sense, such as urine, respiratory tract air, gastrointestinal contents and bile, but because they coexist with the human body in life and death, they needed to form part of the thermal analysis as well. Many symmetrically paired organs, such as long bones and ribs, were presented as individual computational tissue domains. Many tissue domains of smaller body parts such as vitreous humor, intervertebral discs, lens, sclera, cornea, cerebrospinal fluid, blood, cartilage, teeth, tendons/ligaments, diaphragm, dura, mucosa, ureters/urethra, menisci, arteries, veins, spinal cord, lymph nodes, nerves, and so on, were all presented as individual organ domains. Subcutaneous tissue and fat were separate organ domains, and many organs with anatomical subdivisions were divided into separate organ domains, e.g. bone was divided into cancellous, cortical and yellow marrow; cerebrum was divided into white matter, grey matter, midbrain, pons, corpus callosum, hypothalamus, anterior/posterior commissure, pituitary and hippocampus; and kidney was divided into cortex and medulla. All endocrine, exocrine, and genital tissues were represented as separate organ domains (adrenal glands, pituitary gland, pancreas, thyroid gland, prostate gland, salivary gland, epididymis, testis, seminal vesicle, and the penis).

## Methods

### 1. The Finite-Difference Time-Domain Method

The Finite-Difference Time-Domain method (FDTD) is a direct solution of Maxwell’s curl equations in the time domain. It was first proposed by Yee [20] and has since been applied in many fields of engineering, including computational fluid dynamics (CFD). The electric field (E-field) and magnetic field (M-field) components are allocated in space on a staggered mesh of a Cartesian coordinate system. The E-field and H-field components are updated in a leap-frog scheme according to the finite difference form of the curl surrounding the component. The transient field can be calculated when the initial field, boundary, and source conditions are known. Maxwell’s equations are discretized by means of a second-order finite-difference approximation, both in space and in time in an equidistant mesh. For the explicit finite difference scheme to yield a stable mesh solution, the time-step used must be limited according to the Courant-Friedrich-Levy criterion [21]. Time-step is directly related to the size of the grid cell, so that in an equidistantly spaced mesh, a reduction of the mesh step size by a factor of two increases the storage space requirement by a factor of 8 and the computational time by a factor of 16.

### 2. Transient Thermal Analysis Scheme

Transient thermal analysis is a numeric method of solving heat transfer problems over a specified interval. Transient analysis is used to determine either the interval of heat transfer required for temperature equilibration, or to predict temperature distribution after equilibration or after a specified time interval. In this study, transient thermal analysis was used to predict total-body temperature distribution as if the body were alive. This was done by simulating the combined effects of endogenous metabolic heat, heat transfer by blood flow, heat transfer between tissues by thermal conduction, and heat transfer between the skin and ambient air by thermal convection and thermal radiation. The governing equation applied in the transient analysis was the PBE modified to linearize metabolic and blood perfusion rates with temperature, which better mimicked the body’s thermal responses.

#### 2.1 Simulation Setup

The chosen pre-mortem physical activity to be simulated was jogging in the nude on a treadmill at 9 km/h at an ambient temperature of 25°C with no wind. The simulation time was set to 10 000s (2 hours 46 minutes). Density, heat capacitance, thermal conductivity and blood convection temperature values (IT’IS Foundation) were assigned to the 3D model’s 247 organ domains. Jogging at 9 km/h on a flat surface has a total-body metabolic value of 154W [22], therefore metabolic rates of all the 3D model’s thermogenic organs were adjusted to give a sum of 154.06W, and were then kept constant throughout the simulation interval. Metabolic proportional blood flow rates were applied to each organ domain and were also kept constant. The blood convective temperature of each organ, denoted by T_b_ in the PBE, was set to 37°C for all organs except for its skin and subcutaneous tissue, for which it was set to 28°C and 33°C respectively.

#### 2.2 Initial Conditions

Initial conditions refer to the temperature of each organ domain at the beginning of analysis, T_0_. Exact temperature values were not critical for this analysis because they would be determined by the analysis itself irrespective of values given. All organs were assigned initial conditions of 37°C by default.

#### 2.3 Boundary Condition

Boundary conditions specify the thermal conditions at the boundaries of the 3D human model, i.e. skin, using various types of conditions. The three types boundary conditions usually applied in thermal analyses are:

- Dirichlet, in which the surrounding air has a fixed temperature,
- Neumann, in which there is a fixed heat flux present at the interface, with heat, either passing in or out of the body, and
- Mixed, in which the two types coexist.

In a ‘mixed’ boundary condition, the convective heat flux is the product of the heat transfer coefficient *h* and local temperature difference between skin and air temperatures. Mixed boundary condition work well for modelling effects of unforced convection, i.e. in the absence of wind, and is expressed mathematically as:

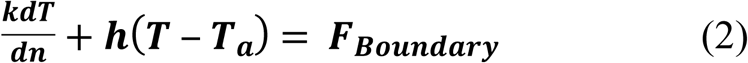

where *T*_*a*_ represents the ambient air temperature. Exchange of radiative heat, derived from the Stefan and Boltzmann equation, in the form:

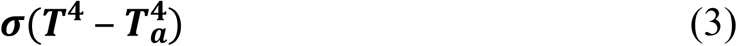

is expressed as:

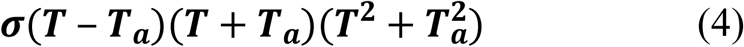

which, for small temperature differences can be approximated to:

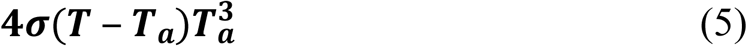

and described as well as a mixed boundary condition with 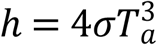. The boundary condition type chosen for this study was mixed: air temperature was 25°C; the total-body heat transfer coefficient (h) was 3.4 Wm-2°C^−1^ [23]; and heat flux was set to 0 W m^−2^.

### 3. Sensors

Sensors are time-points along the analysis interval, where images of the temperature solution are captured for later viewing. Sensors of a chosen number are placed evenly along the simulation interval. Snapshots may later be viewed in quick succession as a video. Thirty-four sensors were placed in the simulation (including at times □ 0 and □ = 10000 □), giving image solutions every ∼300s, or 5 minutes. This was to view temperature evolution from initial conditions to equilibrium.

### 4. Grid Settings

“The numerical solution of partial differential equations such as the PBE requires some discretization of the field into a collection of points or elemental volumes (cells). The differential equations are approximated by a set of algebraic equations on this collection, and this system of algebraic equations is then solved to produce a set of discrete values that approximates the solution of the partial differential system over the field. The discretization of the field requires some organization for the solution thereon to be efficient, i.e. it must be possible to readily identify the points or cells neighboring the computation site. Furthermore, the discretization must conform to the boundaries of the region in such a way that boundary conditions can be accurately represented. This organization is provided by a coordinate system, and the need for alignment with the boundary is reflected in the routine choice of Cartesian coordinates for rectangular regions, cylindrical coordinates for circular regions, etc.” [24].

“A numerically generated grid is understood here to be the organized set of points formed by the intersections of the lines of a boundary-conforming curvilinear coordinate system. The cardinal feature of such a system is that some coordinate line (surface in 3D) is coincident with each segment of the boundary of the physical region. The use of coordinate line intersections to define the grid points provides an organizational structure, which allows all computation to be done on a fixed square grid when the partial differential equations of interest have been transformed so that the curvilinear coordinates replace the Cartesian coordinates as the independent variables” [24]. Sim4Life® automatically constructs a structured grid around any 3D geometry at a click of a button. The number of grid cells can be adjusted by adjusting the maximum step value. The grid size is constructed around our 3D human model was 34.595e+006 cells (142x, 279y and 899z).

### 5. Voxel Settings

A voxel is a discrete volume-pixel that represents a value on a regular grid in three-dimensional space. Each organ domain was divided into multiple voxels whose size and number can be manually adjusted, where a voxel represented a single data point that contained scalar values such as thermal property values, metabolic rates, and blood flow rates as discussed in previous sections. Voxels for all organ domains were created by an automated function of Sim4Life®.

### 6. Solver Settings

The thermal solver applies the conformal mode to alleviate the disadvantageous staircasing effects inherent at surfaces by interpolating to the actual model surface element, but infers increased computational costs. The number of iterations was set at the default maximum value of 1.0e+003 as the iteration number that produced the required relative tolerance was not known in advance. Relative tolerance evaluated convergence and was set at 1.0e–004.

## Results

The transient thermal analyses in this study computed heat transfer between thermogenic and nonthermogenic tissue domains. The latter were stomach lumen, small intestine lumen, large intestine lumen, urine, blood, teeth, vitreous humor, eye lens, eye cornea, cerebrospinal fluid, bile, bronchus lumen, trachea lumen, air, esophagus lumen, and intervertebral discs. Heat transfer by thermal conduction between the treadmill and soles of the feet was neglected. Out of the set 10 000s simulation interval, temperature equilibration occurred at 8400s. False-color thermal mapping of the simulation solution indicated non-uniform but systematic skin temperature distribution that had a strong resemblance to infrared thermal imaging [24] (Fig 2). Skin temperatures ranged from 20.5°C on the bottom of the right heel to 41.7°C on the vertex scalp. Skin temperature distribution demonstrated a strong correlation of body surface contour and to the anatomic location of underlying viscera–– particularly muscle. Temperatures over distal feet and toes, nose tip, anterior and posterior shoulders, and shoulder blades were relatively low, at around 25°C. Higher skin temperatures were on the left axillary fossa and adjacent lateral chest and left arm; on the posterior neck; above the sternal notch; above the clavicles; in both formal regions; on the frontal and parietal scalp; and on the lateral aspects of the forearms. Areas of forearms and the scalp showed temperatures of more than 37.9°C. The pattern of temperature distribution in subcutaneous tissue and muscle was comparable to that of skin.

**Fig 2.**
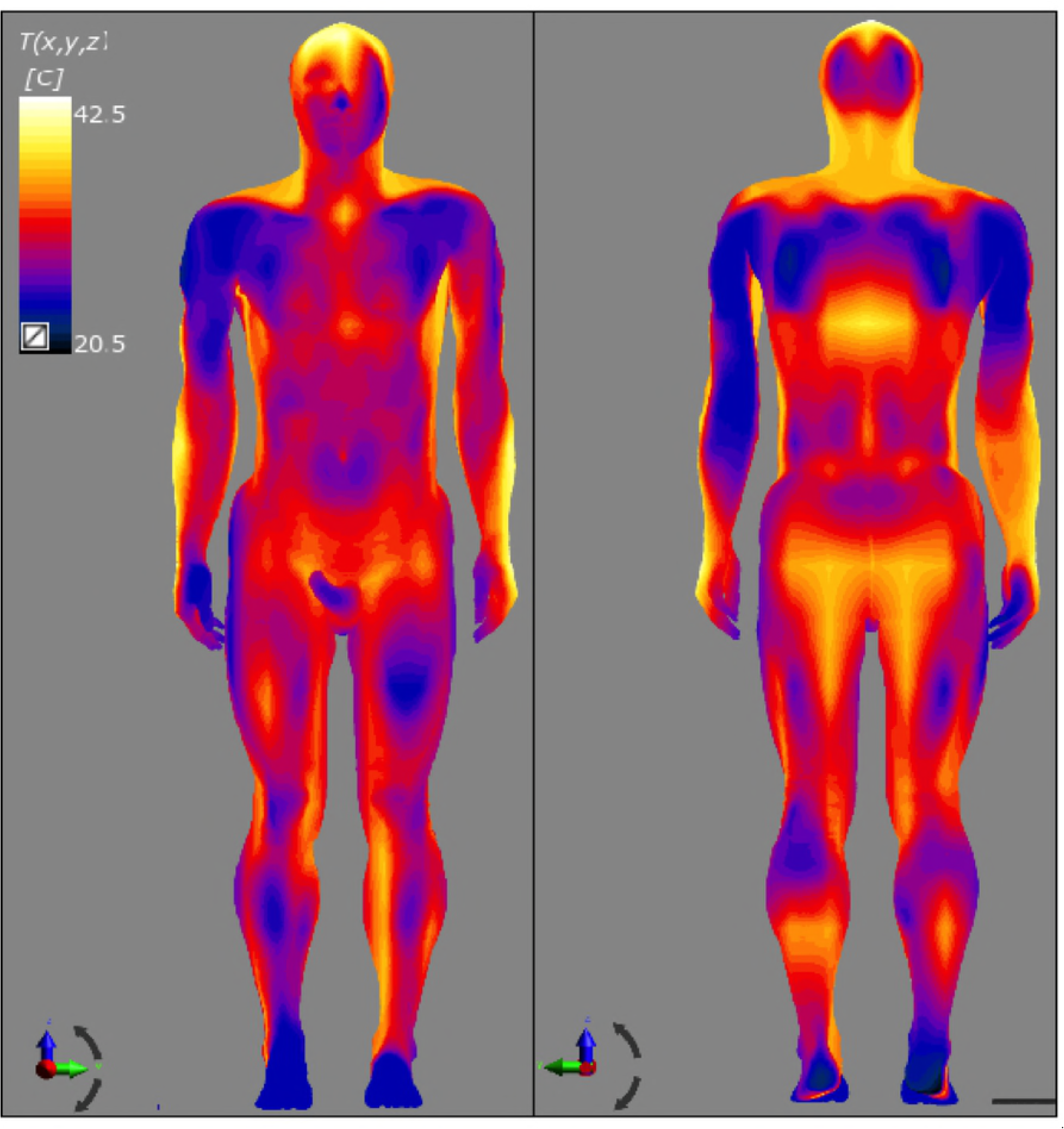
Results of the numerical simulation indicating predicted skin temperature of the 3D human model. On the left is anterior view and on the right is posterior view.

The two regular body sites of deep-core thermometry for death-time calculation using empiric and numeric methods are the deep rectum and liver. The latter is used in instances where rectal thermometry would disturb evidence, e.g. in cases where sexual assault is suspected. We examined predicted temperature of the liver and rectum. In our 3D model, the liver spanned from a height of 1.139m to 1.301m over 81 axial slices. Its anatomic relations were the diaphragm, stomach, small intestines, large intestines, ribs, and muscles. Predicted liver capsule temperatures were topographically inhomogeneous and lower than those of the deep parenchyma (Fig 3). The anterior vertical part of the right liver lobe showed lower capsular temperatures ranging from 32.5°C to 35°C. The capsule on the inward-facing surface revealed higher temperatures compared to the anterior surface–– the highest being 37.6°C. The liver parenchymal temperature was more uniform, with the inferior and inner edge of the left lobe adjacent to the stomach showing a temperature of ±36.1°C, with most of the liver parenchyma averaging at 37.2°C.

**Fig 3.**
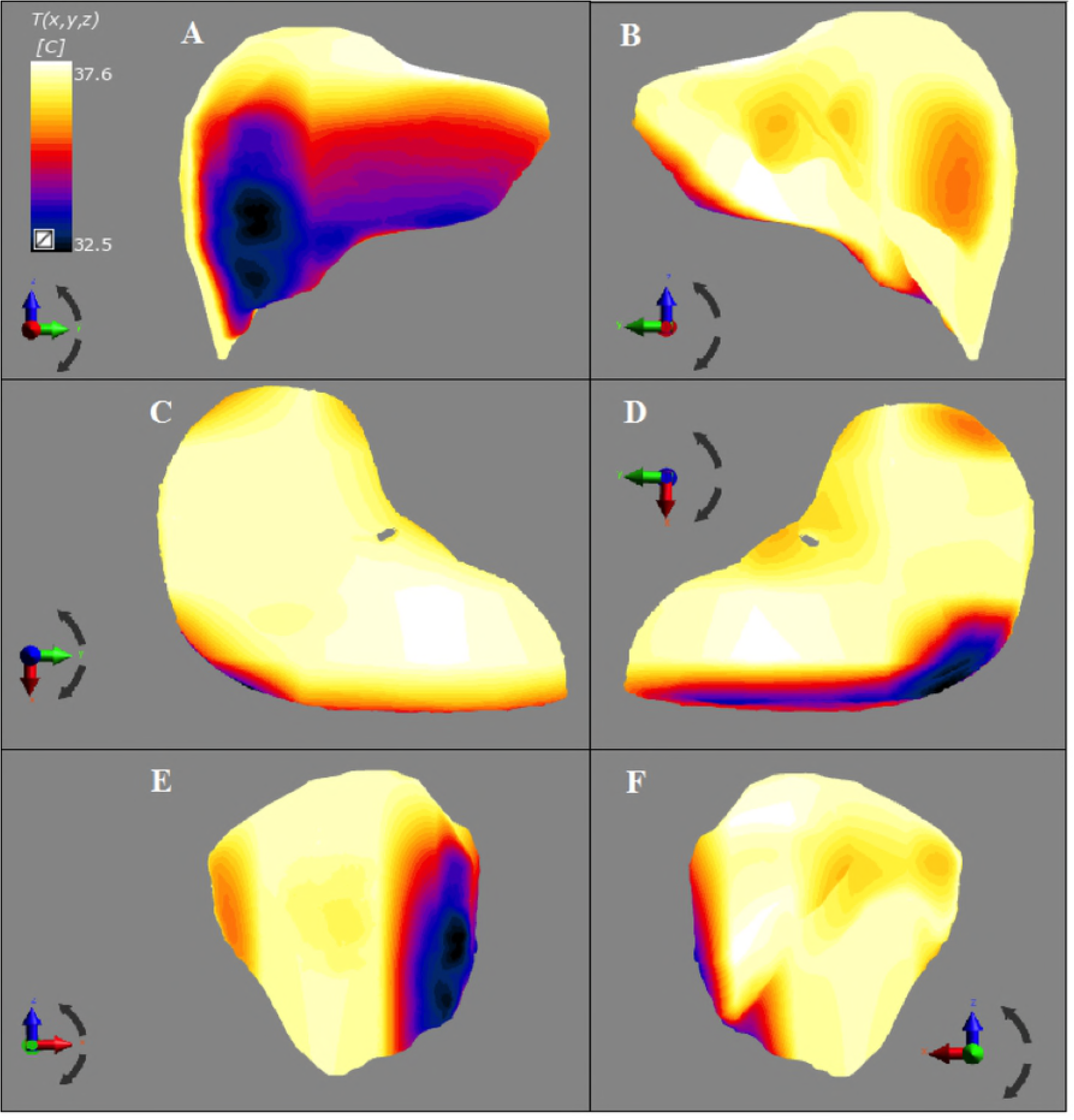
Predicted liver capsule temperature. Figs 3A to 3F show anterior, posterior, superior, inferior and medial views, respectively.

The rectum spanned from a height of 0.83m to 0.96m. Its anatomic relations were the bladder, sacrum and coccyx, iliopsoas muscles, adipose tissue, small intestines and sigmoid colon. The deepest part of the rectum, in which rectal thermometry is performed, showed a predicted temperature of 37.2°C. Axial slices of the 3D human model at both the liver and deep rectal levels (Fig 4) indicated a temperature gradient of ∼3°C between skin and inner core. Predicted surface temperatures of nearly all computational organ domains showed topographic inhomogeneity similar to that of the liver, while their deep parenchymal temperatures were generally higher and more uniform. The length of this paper, however, does not permit for a thorough examination of all organs.

**Fig 4.**
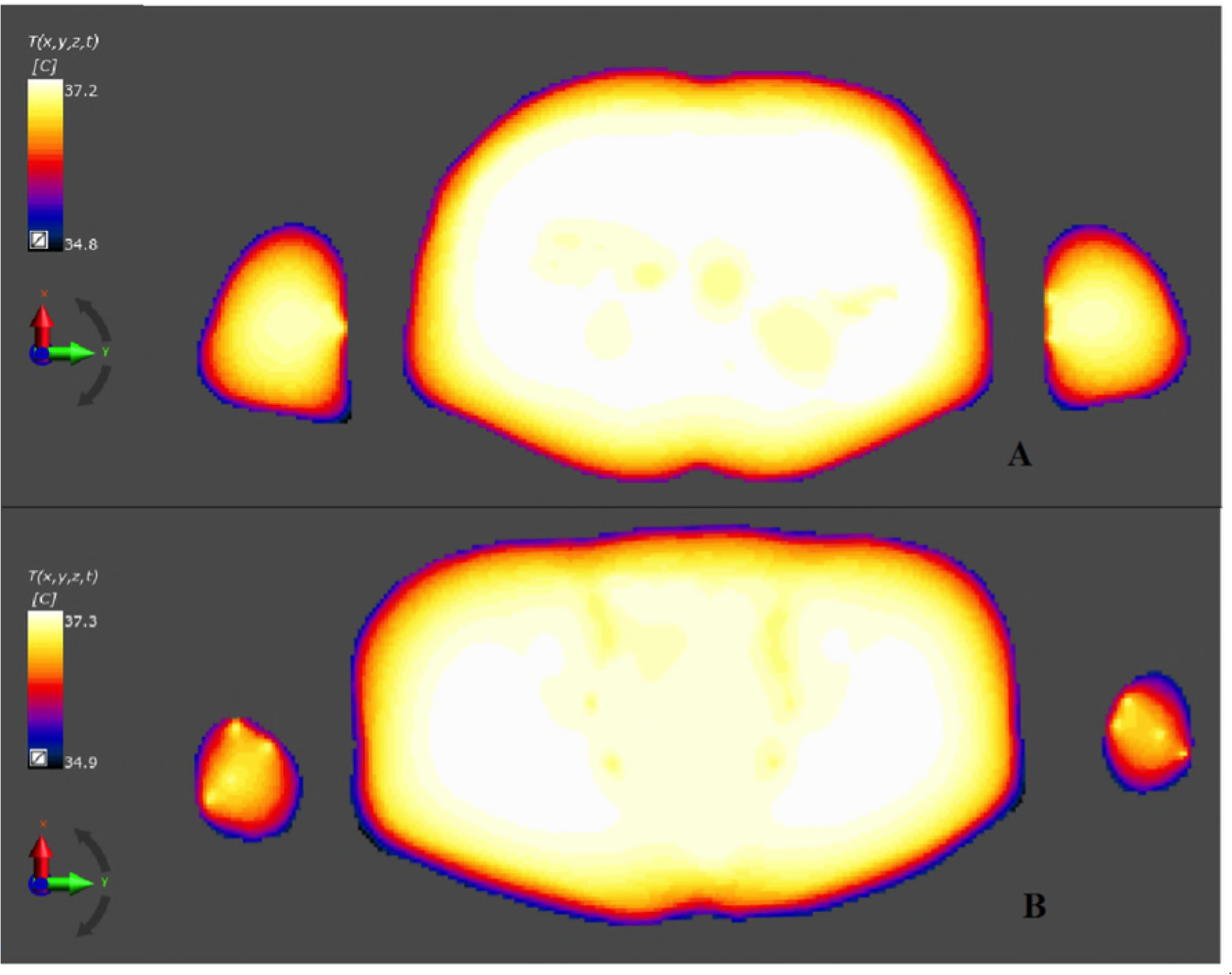
Axial views of predicted body temperature of the male 3D human model. Image A is at the level of the liver and image B is at the level of the deep rectum.

## Discussion

The presumed premortem physical state of a decedent can be determined from death-scene investigation to facilitate adjustment of blood perfusion and metabolic rates in thermonumeric analysis of determining premortem 3D temperature distribution. In life, blood perfusion to each organ is regulated at the minimum level that will supply the tissue’s metabolic needs–– no more, no less [25]. Skeletal muscles, cardiac muscle, lungs, intercostal muscles, the diaphragm, and the adrenal glands are among organs in which metabolic and blood perfusion rates increase during physical exertion. In these organs the blood perfusion rate can be as much as 20 to 30 times the resting level. The heart meets the body’s metabolic demands by increasing its cardiac output, which can increase by no more than four to seven times that of the resting level. The average resting coronary blood perfusion rate is 225ml min^−1^, but increases only threefold to fourfold during strenuous exercise to supply extra nutrients required by heart muscles. The metabolic activity of the skeletal muscle in the resting state is very low. This is so the muscle blood perfusion rate is 4ml 100g ^−1^ min^−1^. During intensive exercise, however, this rate can reach 90ml 100g^−1^ min^−1^ [26]. Blood perfusion through the lungs is equal to cardiac output, and is therefore influenced by the same factors. Metabolic heat generation and blood perfusion rates of intercostal muscles and the diaphragm are regulated to meet respiratory efforts associated with exertion. The software used in this study allowed only manual adjustment of these parameters, and an automated function would improve efficiency.

Thermonumeric analysis for calculating premortem 3D temperature distribution ideally ought to include a system that mimics the body’s thermoregulatory system, which consists of hot and cold temperature receptors in the skin, anterior hypothalamus, and deep viscera. Afferent temperature signals terminate at the anterior hypothalamus and efferent thermoregulatory impulses originate from the posterior hypothalamus. Together the anterior and posterior hypothalamus maintain a core body temperature ranging from 36°C to 37.5°C. Skin temperature fluctuates according to ambient temperatures, clothing, and the state of thermoregulation. The critical anterior hypothalamic temperature set-point beyond which any thermoregulatory responses are activated is 37.1°C [25]. Mechanisms to increase body temperature are cutaneous vasoconstriction, inhibition of sweating, muscle tensioning in the skin, non-shivering (chemical) thermogenesis, shivering, and thyroxin secretion. The anterior hypothalamic temperature set-points for vasoconstriction and shivering are 37°C and 35.9°C, respectively. These set-points are raised during exercise and fever. Cutaneous vasoconstriction reduces heat transfer from blood to skin, and blood supply to the cutaneous vascular plexus can be effectively as low as zero, resulting in a local temperature gradient across a skin temperature of 10°C [27]. Such vasoconstriction causes ischaemia and eventually gangrene when prolonged, e.g. frostbite in extreme cold. Shivering is the body’s mechanism to increase temperature and can be initiated by cooler skin despite one’s hypothalamus temperature being above 37.1°C. During maximum shivering, body heat generation can rise to four to five times the normal basal metabolic heat generation rate [25]. The hypothalamic set-points for both shivering and vasoconstriction are raised during physical exertion, so that shivering does not occur during exercise in cold weather and when the skin is cold but a person’s core is hot. Shivering stops when the core temperature falls below 30°C [28]. Mechanisms to decrease body temperature involve cutaneous vasodilation, sweating, and a decrease in metabolic heat generation. The anterior hypothalamic temperature set-point for both cutaneous vasodilation and sweating is 37.2°C [27]. The set-point for sweating is altered by cutaneous temperature receptors so that at the same core temperature warmer skin enhances sweating and a cooler skin inhibits it [16, 27]. Sweating is the secretion of salty water onto the skin by eccrine sweat glands controlled by the autonomic nervous system. Sweat is evaporated by heat transferred from skin, resulting in skin cooling, and therefore acts as a heat sink and increases heat flux out of the body. Maximum sweating rates of 2 to 3L/hour can remove heat from the body at a rate of more than 10-fold the normal basal metabolic heat generation. Sweating increases the heat transfer coefficient of skin by up to 30-fold compared to dry skin [29]. Although individual blood perfusion and metabolic and rates of the organs in the 3D human model could theoretically be adjusted manually, the method used did not have a thermoregulatory system, which would have needed to be designed and programmed from scratch. Shivering and sweating were thus not represented in this study.

## Conclusion

Premortem three-dimensional temperature distribution can be predicted accurately using stand-alone transient thermonumeric analysis, and high-end 3D human models developed from MRIs of a healthy, living adult subjects are the best human surrogates for the purpose. Organ-specific metabolic heat and blood flow rates can be adjusted for a premortem physical activity judged by a death-scene investigation. This paper advocates for separation of thermonumeric analysis for determining premortem three-dimensional temperature distribution from thermonumeric analysis representing postmortem cooling. The separation clarifies the transition from life to death, which is marked by a cessation of organ blood flow and metabolism. The proposal is that forensic pathologists should estimate the pre-mortem physical activity of a decedent using findings from death-scene investigation, after which a corresponding metabolic value of that activity should be established from the literature. Organ metabolic and blood flow rates should then be adjusted accordingly when performing a thermonumeric analysis to determine premortem 3D temperature distribution.

Thermoregulation for the kind of 3D human models used in this study can be improved to the same level of realistic complexity as those used in HVAC systems as discussed before. The convective blood temperature of each organ domain, which in the study was set to 37°C, ought to be solved for as part of the transient analysis rather than being prescribed by the user or presumed to be universally and continuously constant. Unfortunately, the governing PBE in the thermal solver used in this study could not be edited and would have required a new code to be developed from scratch. Explicit solving of blood flow velocity and blood heat flux within the cardiovascular system using CFD would have been ideal as it would have intrinsically determined the convective blood temperature. The computational cost, however, would have been prohibitively expensive. This study analyzed heat transfer of a naked body only, and clothing simulation was beyond the scope of this paper and may be the subject of future study.

## References

[1] Henssge C (1988). Death Time Estimation in case work I. The rectal temperature time of death nomogram. Forensic Science International, 209–236.

[2] Mall G, Eienmenger W. (2005). Estimation of time since death by heat-flow Finite-Element model. Part I: method, model, calibration and validation. Legal Medicine, 1–14.

[3] Marshall TK, Hoare FE (1962). Estimating the time of death— the rectal cooling after death and its mathematical expression. J Forensic Sci, 56–81.

[4] Marshall TK, Hoare FE (1962). Estimating the time of death— the use of body temperature in estimating the time of death. J Forensic Sci, 11–21.

[5] Henssge C (1979). Die Praszision von Todeszeitschätzungen durch die mathematische Beschreibung der rektalen Leichenabkuhlung. Z Rechtsmed, 49–67.

[6] Althaus L, Zur Todeszeitbestimmung aus der Leichenabkühlung bei sprunghaft wechselnder Umgebungstemperatur, Master’s Thesis, Universitat Essen, 1997.

[7] Althaus L, Henßge C (1999) Rectal temperature time of death nomogram: sudden change of ambient temperature, Forensic Science International 99 171–178.

[8] Smart J (2010). Estimating time of death with a Fourier Series Unsteady-state Heat Transfer Model. Journal of Forensic Science 1481–1487.

[9] Schenkl S, Muggenthaler H, Hubig M, Erdmann B, Weiser M, Zachow S, Heinrich A, Güttler FV, Teichgräber U, Mall G (2017). Automatic CT-based finite element model generation for temperature-based death time estimation: feasibility study and sensitivity analysis, International Journal of Legal Medicine 131:699–712.

[10] Pennes HH (1948). Analysis of tissue and arterial temperatures in the resting human forearm. Journal of Applied Physiology, 93–122.

[11] Atkins AR (1973). Application of the hopscotch algorithm for solving the heat flow equation for the human body. Computational Biology and Medicine, 397–405.

[12] Arkin H, Shitzer A (1984). A model of thermoregulation in the human body. ASME 84-WA/HT-66, 1–7.

[13] Crosbie RJ, Hardy JD, Fessenden E (1961). Electrical Analog Simulation of Temperature Regulation in Man. IRE Transactions on bio-medical electronics, 245–252.

[14] Fiala D, Lomas KJ, Stohrer M (1999). A computer model of human thermoregulation for a wide range of environmental conditions: the passive system. Journal of Applied Physiology, 1957–1972.

[15] Gordon RG, Roemer RB, Horvath SM (1976). A mathematical model of the human temperature regulatory system - transient cold exposure response. IEEE Tran Biomed Eng, 434–444.

[16] Stolwjk AJ (1971). A mathematical model of physiological temperature regulation in man. Springfield, vA: NASA CR-1855.

[17] Wissler EH (1961). Steady-state temperature distribution in man. Journal of Applied Physiology, 734–740.

[18] Wissler, EH (1964). A mathematical model of the human thermal system. Bulletin of Mathematical Biology, 147–165.

[19] Gosselin M, Neufeld E, Moser H, Huber E, Farcito S, Gerber L, Jedensjö M, Hilber I, Di Gennaro F, Lloyd B, Cherubini E, Szvzerba D, Kainz W, Kuster N (2014). Development of a new generation of high-resolution anatomical models for medical device evaluation: the Virtual Population 3.0. Physics in Medicine and Biology, 5287 - 5303.

[20] Yee K (1966). Numerical Solution of initial boundary value problems involving Maxwell’s equation in isotropic media. IEEE Transactions on Antennas and Propagation, 302–207.

[21] Courant R, Friedrichs K, Lewy H (1928). “Über die partiellen Differenzengleichungen der mathematischen Physik” Mathematische Annalen (German), 100 (1): 32–74.

[22] Jette M, Sidney K, Blumchen G (1990). Metabolic Equivalents (METS) in Exercise Testing, Exercise Prescription and Evaluation of Functional Capacity. Clinical Cardiology, 555–565.

[23] de Dear R, Arens E, Zhang H, & et al. (1997). Convective and radiative heat transfer coefficients for individual human body segments. Int J Biometeorol, 141–156.

[24] Costa C, Sillero-Quintana MI, Piñonosa Cano S, Moreira DG, Brito C, Fernandes AJA, Azumbuja Pussieldi G, Marins JCB (2016). Daily oscillations of skin temperature in military personnel using thermography. Journal of the Royal Army Medical Corps, 335–342.

[25] Hall JE, Guyton AC (2006) Textbook of Medical Physiology. Saunders/Elsevier. Philadelphia.

[26] Saltin B, Radegran G, Koskolou M, Roach RC (1998). Skeletal muscle blood flow in human and its regulation during exercise. Acta Physiologica Scandinavica 421–436.

[27] Arens E, Zhang H (2006). The skin’s role in human thermoregulation and comfort. In N. Pan & P. Gibson, Thermal and Moisture Transport in Fibrous Materials (p. 560). Woodhead Publishing.

[28] Lloyd E (1996). Accidental hypothermia. Resuscitation, 111–124.

[29] Alber-Wallerstrom B, Holmer I (1985). Efficiency of sweat evaporation in unacclimatized man working in a hot humid environment. Eur J Appl Physiol, 480–487.

